# Dissecting and steering cell dynamics using spatially-informed RNA velocity with veloAgent

**DOI:** 10.64898/2026.01.09.698589

**Authors:** Brent Yoon, Vishvak Raghavan, Jun Ding

## Abstract

RNA velocity enables inference of cell state transitions from single-cell transcriptomics by modeling transcriptional dynamics from spliced and unspliced mRNA. However, existing methods overlook spatial context and struggle to scale to large datasets, limiting insights into tissue organization and dynamic processes. We introduce veloAgent, a deep generative and agent-based framework that estimates gene- and cell-specific transcriptional kinetics while integrating spatial information through agent-based simulations of local microenvironments. By leveraging both molecular and spatial cues, veloAgent improves velocity accuracy and achieves sublinear memory scaling, enabling efficient analysis of large and multi-batch spatial datasets. A distinctive feature of veloAgent is its in silico perturbation module, which allows targeted manipulation of spatial velocity vectors to simulate regulatory interventions and predict their impact on cell fate dynamics. These capabilities position veloAgent as a scalable and versatile framework for dissecting spatially resolved cellular dynamics and guiding cell fate manipulation across diverse biological processes.

## MAIN

Dynamic changes in gene expression, including both transcriptional and post-transcriptional regulation are fundamental to determining a cell’s identity and function^1,2^. Single-cell RNA sequencing (scRNA-seq) has revolutionized the field of transcriptomics by enabling high-resolution profiling of gene expression at the single-cell level, thereby facilitating unprecedented insights into cellular heterogeneity, gene regulatory dynamics, and developmental trajectories^3,4^. In parallel, advances in spatial transcriptomics have made it possible to preserve spatial context while quantifying gene expression, enabling researchers to investigate tissue organization, niche-specific expression programs and cell-cell interactions^5–7^. Despite its quantitative precision and comprehensive coverage, these technologies only provide a temporally static snapshot of cellular states rather than a trajectory of cellular behaviour. This limitation constrains our ability to reconstruct dynamic processes such as differentiation, and lineage commitment^8^. Addressing these limitations, RNA velocity has emerged as a transformative framework for inferring future cellular states by leveraging the relative abundances of unspliced and spliced mRNA transcripts to model ongoing gene expression kinetics^9,10^. This approach not only enables the extrapolation of transcriptional dynamics and trajectory prediction but also offers critical insights into cellular decision-making processes with broad applications in developmental biology, and disease progression.

Several computational frameworks have been developed to infer RNA velocity, including cellDancer^11^, scVelo^10^, SIRV^12^, DeepVelo^13^, VeloVAE^14^, VeloVI^15^, and UniTVelo^16^. These methods introduce various strategies to improve accuracy, such as incorporating dynamical systems modeling, deep generative modeling, and kinetic parameter estimation. However, existing RNA velocity methods face several limitations that constrain our ability to fully understand development and disease progression. Many existing methods still assume uniform transcriptional kinetics across genes or cells, an oversimplification that erases the cell-to-cell heterogeneity that drives development, disease onset and progression. While more advanced models such as cellDancer estimate gene- and cell-specific kinetics and yield biologically plausible results, their heavy computational demands make them impractical for the large, multi-batch datasets increasingly central to development or disease atlas-scale studies¹¹. Perhaps, most critically, nearly all current approaches ignore spatial context, relying instead on embeddings of transcriptional similarity. This omission ignores vital information from tissue microenvironments and cell–cell interactions, leading to inferred trajectories that cross anatomical boundaries or contradict known tissue structure^17–19^. This loss of spatial fidelity prevents us from uncovering how tumors invade, how immune cells coordinate in tissue, or how stem cells rebuild organs^20–22^. Furthermore, no current framework allows researchers to computationally perturb key genes or pathways and predict how such perturbations reshape cellular trajectories. Such a tool would be especially powerful in regenerative medicine and disease intervention, where rapid in silico tests could help prioritize experiments, highlight therapeutic targets, and significantly reduce the time and cost of discovery pipelines. Together, these gaps limit not just technical accuracy but our ability to translate single-cell maps into insights about developmental and disease biology. This limitation has also constrained our understanding of how cellular dynamics unfold within the structural and spatial context of native tissues.

To address these challenges, we introduce veloAgent, a hybrid framework consisting of three distinct components: a variational autoencoder (VAE), a gene interaction-informed neural network, and an agent-based model (ABM) to infer spatial RNA velocities. The VAE learns robust latent representations from noisy spliced and unspliced mRNA counts, providing a unified and denoised embedding of transcriptional states. The neural network guided by gene-gene interaction networks derived from the STRING database encodes cell- and gene-specific transcriptional kinetics, while operating in a fixed-dimensional, gene-centric space that scales efficiently to a large number of cells^23^. The ABM then leverages spatial transcriptomics data to simulate each cell as an autonomous agent positioned in its native tissue context, whose future state is influenced by neighbouring cells within its local environment. ABMs have previously used in spatial transcriptomics to model interactions between cells, supporting our choice of this modeling paradigm^24^. This integrative design enables veloAgent to model spatial RNA velocity, capturing how gene expression dynamics unfold across both time and space. By embedding tissue architecture directly into the inference process, we bridge the divide between molecular kinetics and histological organization, transforming RNA velocity from an abstract representation into a spatially coherent model of cell-state transitions.

We demonstrate the utility of veloAgent across a range of spatial transcriptomics platforms with varying spatial resolutions, including Visium, Stereo-seq, HybISS, and seqFISH^25–28^. Unlike existing RNA velocity methods, veloAgent does not simply infer trajectories from just the transcriptome, but directly integrates spatial coordinates, enabling analysis of velocity vectors onto tissue architecture. This uniquely allows researchers to resolve dynamic cellular programs within native niches, revealing transition pathways that respect anatomical boundaries and microenvironmental organization rather than producing over-smoothed, anatomically implausible fields. veloAgent is also designed for scale with its gene-centric framework that makes velocity analysis practical on atlas-scale and multi-batch datasets. Finally, veloAgent introduces an innovation not previously available in velocity analysis: in silico perturbation of transcriptional kinetics. This enabling virtual screening and prioritization of interventions, generating testable hypotheses that reduce experimental cost and accelerate translational discovery. By integrating transcriptomic kinetics, gene-network priors, and spatial context, veloAgent enables both mechanistic interpretation and in-silico perturbation, defining a new paradigm for data-driven hypothesis generation and therapeutic target discovery.

## RESULTS

### Overview of veloAgent

veloAgent introduces a hybrid deep learning and agent-based modeling (ABM) framework for estimating RNA velocity at single-cell resolution by integrating gene expression data with prior gene–gene interaction networks and spatial information. Unlike existing RNA velocity frameworks, veloAgent jointly models transcriptional kinetics and spatial context within a unified deep generative–agent hybrid. As illustrated in Fig. 1, the methodology proceeds through four main stages. First (Fig. 1a), a variational autoencoder (VAE) encodes each cell’s spliced and unspliced mRNA counts into a unified latent representation. This step reduces dimensionality while preserving key transcriptional features that reflect dynamic cellular states. Second (Fig. 1b), the resulting latent vectors are processed through a biologically constrained gene–gene deep neural network (DNN) to infer cell- and gene-specific kinetic parameters—transcription rate (*α*), splicing rate (*β*), and degradation rate (*γ*). The network architecture is guided by protein–protein interactions from the STRING database, enforcing biologically grounded sparsity: each gene’s output depends only on its annotated interactors. This design enables the model to capture meaningful regulatory dependencies during kinetic inference. The inferred parameters are then incorporated into differential equations to compute initial RNA velocity estimates.

**Fig. 1:**
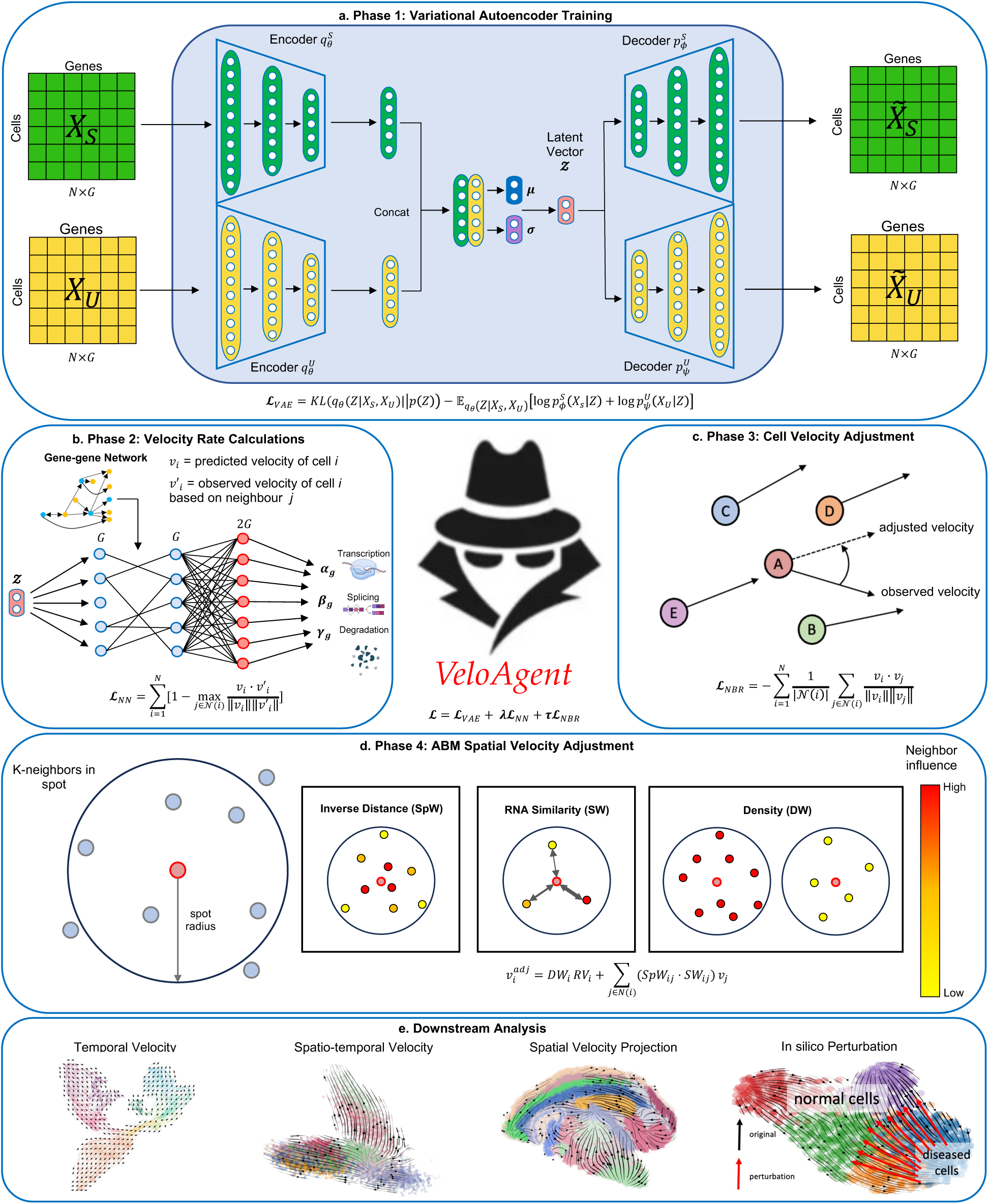
Overview of the veloAgent framework. veloAgent is a hybrid deep learning and agent-based modeling framework for estimating RNA velocity and decoding cellular dynamics by integrating spliced and unspliced transcriptomics with spatial information. The model operates in four main phases, followed by versatile downstream applications. **a, Phase 1: Latent representation learning.** Spliced and unspliced gene expression matrices are separately encoded using variational autoencoders (VAEs). The resulting embeddings are concatenated into a shared latent space *Z*, which captures underlying cellular states while minimizing reconstruction error. **b, Phase 2: Cell- and gene-specific kinetic rate inference.** Kinetic parameters—transcription (*α*), splicing (*β*), and degradation (γ) rates—are inferred for each gene in each cell by passing the latent vector *Z* through a gene-informed neural network. The network is trained to maximize cosine similarity between predicted and observed RNA velocity vectors across cells. **c, Phase 3: Velocity vector refinement.** Cell-level velocity vectors are refined to enforce local coherence in the latent space, aligning similar cells toward consistent velocity directions and improving trajectory continuity. **d, Phase 4: Spatial RNA velocity refinement.** When spatial transcriptomics data are available, an agent-based model further refines velocity vectors by incorporating spatial priors, weighting the influence of neighboring cells based on intercellular distance and expression similarity. **e, Downstream applications.** veloAgent supports multiple downstream analyses, including temporal and spatially informed velocity estimation, projection of velocity fields onto spatial tissue maps, and in silico perturbation experiments to identify candidate interventions that may reverse disease-associated cell state transitions.

The third stage (Fig. 1c) refines the estimated velocity field by enforcing local coherence. Each cell’s velocity vector is aligned with those of its transcriptionally similar neighbors, minimizing directional noise and enhancing trajectory continuity. Finally (Fig. 1d), for datasets with spatial transcriptomics measurements, veloAgent integrates an ABM to further refine velocities in their spatial context. The ABM incorporates intercellular distance, local cell density, and expression similarity to simulate spatial constraints, yielding biologically realistic and spatially informed velocity fields in structured tissue environments. Beyond velocity inference, veloAgent provides multiple outputs for downstream analysis—including temporal and spatial velocity projections—and features a perturbation module that enables in silico steering of RNA velocity directions to model potential gene perturbations (Fig. 1e). Together, these components establish veloAgent as a unified framework for reconstructing and perturbing cellular dynamics in both temporal and spatial contexts.

### veloAgent enables spatially informed RNA velocity inference

veloAgent extends RNA velocity analysis by embedding spatial information directly into the inference process, allowing trajectories to better reflect tissue architecture and neighborhood organization. Through an agent-based spatial refinement module, the veloAgent integrates intercellular proximity, local density, and expression similarity to couple neighboring cells during velocity estimation. This spatial coupling promotes locally aligned velocity directions and suppresses noise arising from isolated or contradictory cell states. As a result, veloAgent produces coherent and biologically interpretable velocity fields that conform to underlying tissue structure and resolve directional ambiguities commonly observed in conventional methods.

To evaluate its performance, we applied veloAgent to four representative spatial transcriptomics datasets—mouse brain (Stereo-seq), breast cancer, chicken heart, and mouse brain (HybISS). Across all datasets, veloAgent achieved superior performance in both qualitative and quantitative benchmarks, demonstrating that spatial context provides a more faithful representation of tissue-level dynamics. In most regions, veloAgent and SIRV yielded broadly consistent trends; however, veloAgent clearly outperformed in complex or transitional zones where spatial organization was critical. In the Stereo-seq mouse brain, for example, SIRV produced unclear or contradictory velocities around fiber-tract cells, whereas veloAgent captured coherent transitions from hippocampus, thalamus sensory-motor (DORsm), and polymodal association (DORpm) regions, consistent with known projections of fiber-tract neurons (Fig. 2a)^29–32^. In breast cancer, SIRV incorrectly directed flows from tumor cells toward normal cells, whereas veloAgent corrected these mispredictions, yielding biologically consistent trajectories (Fig. 2b)^33^. In the chicken heart, veloAgent corrected SIRV’s reversed trajectories at the ventricle-atrium boundary and valve cluster (Fig. 2c)^34^. Similarly, in the HybISS brain, SIRV misdirected neuron velocities toward neuroblasts, whereas veloAgent correctly modeled neurons as terminal states (Fig. 2d)^35^. Collectively, these results highlighted veloAgent’s ability to recover biologically consistent transitions where other approaches failed.

**Fig. 2:**
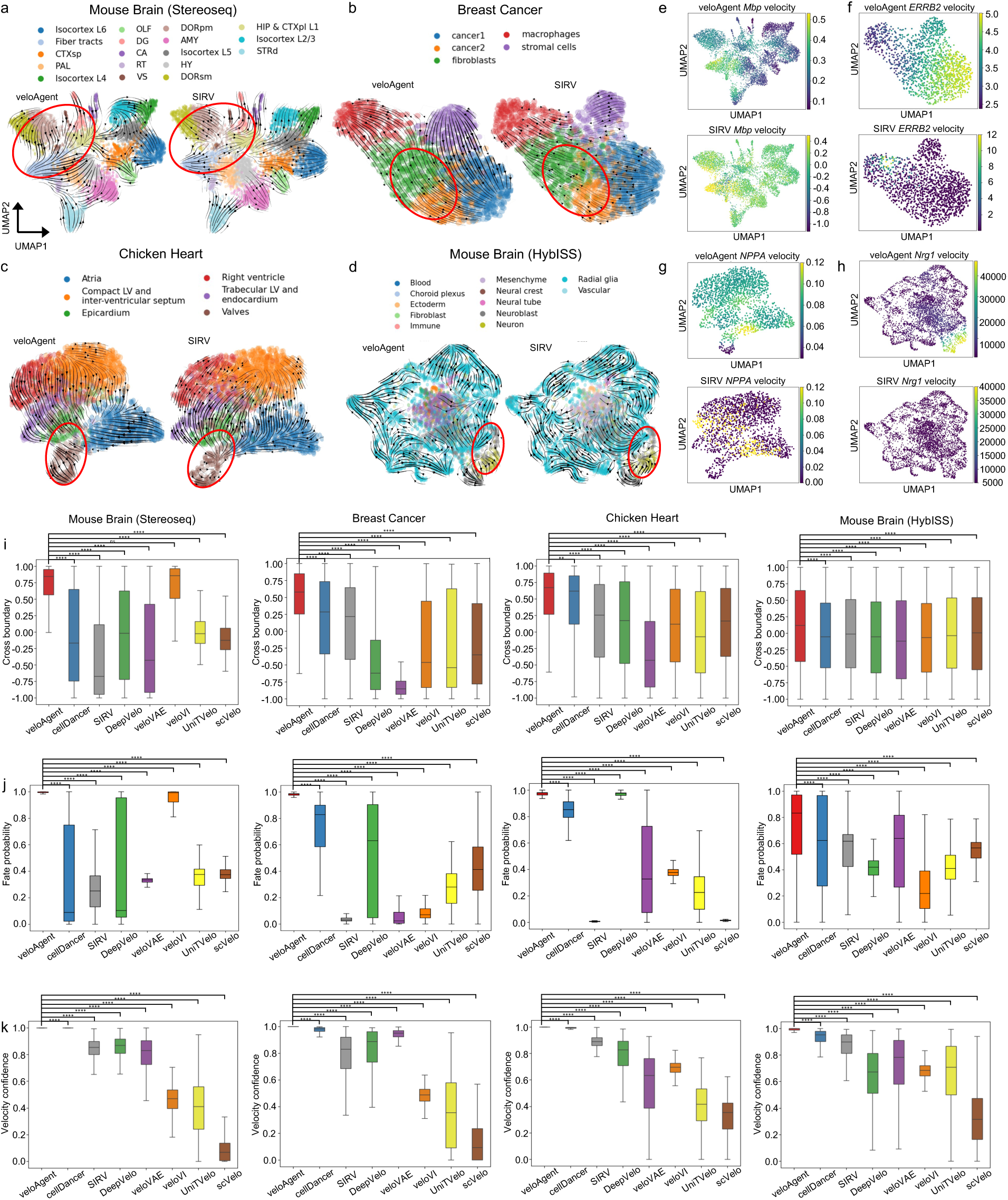
Performance benchmarking of veloAgent for spatially informed RNA velocity inference. This figure evaluates veloAgent’s ability to infer RNA velocity in spatial transcriptomics datasets, comparing its performance to existing methods using both qualitative and quantitative benchmarks. **A-d**, RNA velocity vectors computed by veloAgent (left) and SIRV (right), projected onto UMAP embeddings of four developmental datasets: **a,** mouse brain (Stereo-seq); **b,** breast cancer; **c,** chicken heart; and **d,** mouse brain (HybISS). Red circles highlight key regions of disagreement between methods. **E-h,** Velocity fields of marker genes expressed in terminal cell populations, comparing veloAgent (top) to SIRV (bottom): **e,** *Mbp* in mouse brain (Stereo-seq); **f,** *ERBB2* in breast cancer; **g,** *NPPA* in chicken heart; **h,** *Nrg1* in mouse brain (HybISS). veloAgent demonstrates stronger alignment with expected terminal regions. **i,** Cross boundary scores across benchmarked methods (veloAgent, cellDancer, SIRV, DeepVelo, veloVAE, veloVI, UniTVelo, scVelo), measuring the correctness of predicted transitions across annotated cluster boundaries, based on ground-truth developmental labels. Higher scores reflect better agreement with known trajectories **j,** Fate probability scores across methods, quantifies the likelihood that a cell will progress toward a defined terminal state based on inferred RNA velocity. **k,** Velocity confidence scores across benchmarked methods, quantifying how well predicted velocities align with local gene expression structure. Higher scores indicate more reliable predictions. Statistical significance calculated using Mann-Whitney U test with FDR correction. Exact *P*-values can be found in Supplementary Table 1.

We next validated the spatial fidelity of inferred velocities using well-established regional marker genes that define terminal or tissue-specific populations (Fig. 2e-h). In each dataset, veloAgent’s velocity fields aligned closely with the known spatial domains of these markers, whereas SIRV frequently produced diffuse or misplaced activity. In the Stereo-seq mouse brain, *Mbp* velocity was confined to fiber tracts, consistent with its role in oligodendrocyte maturation and myelination (Fig. 2e)^35^. In breast cancer, *ERBB2* velocity localized correctly within tumor regions but was erroneously assigned to macrophages by SIRV, contradicting its established role as an oncogene and therapeutic target (Fig. 2f)^36^. In the chicken heart, *NPPA* velocity was restricted to atria under veloAgent yet aberrantly extended into ventricles with SIRV, inconsistent with its function as an atrial differentiation marker distinguishing atria from ventricles (Fig. 2g)^37^. In the HybISS mouse brain, *Nrg1* velocity was specifically confined to neurons, consistent with its role in neuronal development, while SIRV failed to capture any localized signal (Fig. 2h)^38^. These spatial-marker-based evaluations confirm that veloAgent faithfully recapitulates known anatomical domains and improves the spatial interpretability of RNA velocity.

Quantitative benchmarking against seven leading RNA-velocity models (cellDancer, SIRV, DeepVelo, veloVAE, veloVI, UniTVelo, scVelo) confirmed these gains (Fig. 2i-k). Evaluated across three complementary metrics—fate probability^39^, cross-boundary direction correctness^40^, and velocity confidence^40^—veloAgent achieved the highest or near-highest scores in all datasets. For cross-boundary direction correctness, veloAgent was on average 92.03% (Stereo-seq brain), 92.02% (breast cancer), 55.52% (chicken heart), and 12.15% (HybISS brain) better than all other methods. Specifically, veloAgent matched veloVI in Stereo-seq brain (0.8447 vs. 0.8590, n.s) and outperformed all methods in the other datasets: breast cancer (0.6148 vs. 0.2849, cellDancer; FDR = 3.588×10^-60^), chicken heart (0.6742 vs. 0.6193, cellDancer; FDR = 2.923×10^-3^), and HybISS brain (0.1237 vs. –0.0209, SIRV; FDR = 3.880×10^-3^). For fate probability, veloAgent consistently showed stronger or comparable predictive power in comparison to other methods and on average, veloAgent achieved 64.97% (Stereo-seq brain), 65.87% (breast cancer), 58.12% (chicken heart), and 33.32% (HybISS brain) higher predictive power than all competing methods. Specifically, veloAgent consistently demonstrated stronger or comparable performance: Stereo-seq brain (0.9995 vs. 0.9919, veloVI; FDR = 6.990×10^-132^), breast cancer (0.9841 vs. 0.6309, DeepVelo; FDR < 1.0×10^-308^), chicken heart (0.9779 vs. 0.8520, cellDancer; FDR = < 1.0×10^-308^), and HybISS brain (0.8324 vs. 0.6231, cellDancer; FDR = 1.071×10^-51^). For velocity confidence, veloAgent achieved on average 35.61% (Stereo-seq brain), 34.92% (breast cancer), 31.23% (chicken heart), and 28.25% (HybISS brain) higher internal consistency than all competing methods. Specifically, veloAgent maintained near-perfect internal consistency (≥ 0.998) across datasets: Stereo-seq brain (0.9995 vs. 0. 8704, DeepVelo; FDR < 1.0×10^-308^), breast cancer (0.9987 vs. 0.8304, SIRV; FDR < 1.0×10^-308^), chicken heart (0.9999 vs. 0.6349, veloVAE; FDR < 1.0×10^-308^), and HybISS brain (0.9987 vs. 0.9524, cellDancer; FDR < 1.0×10^-308^). These improvements reflect more accurate modeling of transition boundaries, terminal fates, and noise-robust velocity estimation.

Although principally designed for spatial data, veloAgent also applies to conventional scRNA-seq datasets, maintaining advantages over existing approaches (Supplementary Figs. 1-2). Extended benchmarking across additional datasets and metrics (Supplementary Figs. 3-5) further demonstrate its robustness, positioning veloAgent as a powerful framework for reconstructing cellular dynamics with spatial fidelity and quantitative precision.

### Spatially informed RNA velocity reveals tissue-coherent cell-state transitions

Most existing RNA velocity methods infer transcriptional dynamics solely in latent transcriptomic space, treating velocity as an abstract trajectory independent of tissue structure. In contrast, veloAgent integrates spatial priors into RNA velocity inference through an ABM, embedding velocity vectors directly onto tissue coordinates to capture how cellular transitions are shaped by native architecture. This integration enables spatially coherent and biologically grounded reconstruction of cell-state dynamics. Across four spatial transcriptomics datasets, veloAgent produced velocity fields that were directionally coherent, spatially continuous, and extended along broader developmental transition tracks than competing methods (Fig. 3a-d, Supplementary Fig. 6). By enforcing local spatial coherence through ABM simulations, veloAgent revealed how tissue organization constrains and guides cellular trajectories—an effect absent from models that neglect spatial context.

**Fig. 3:**
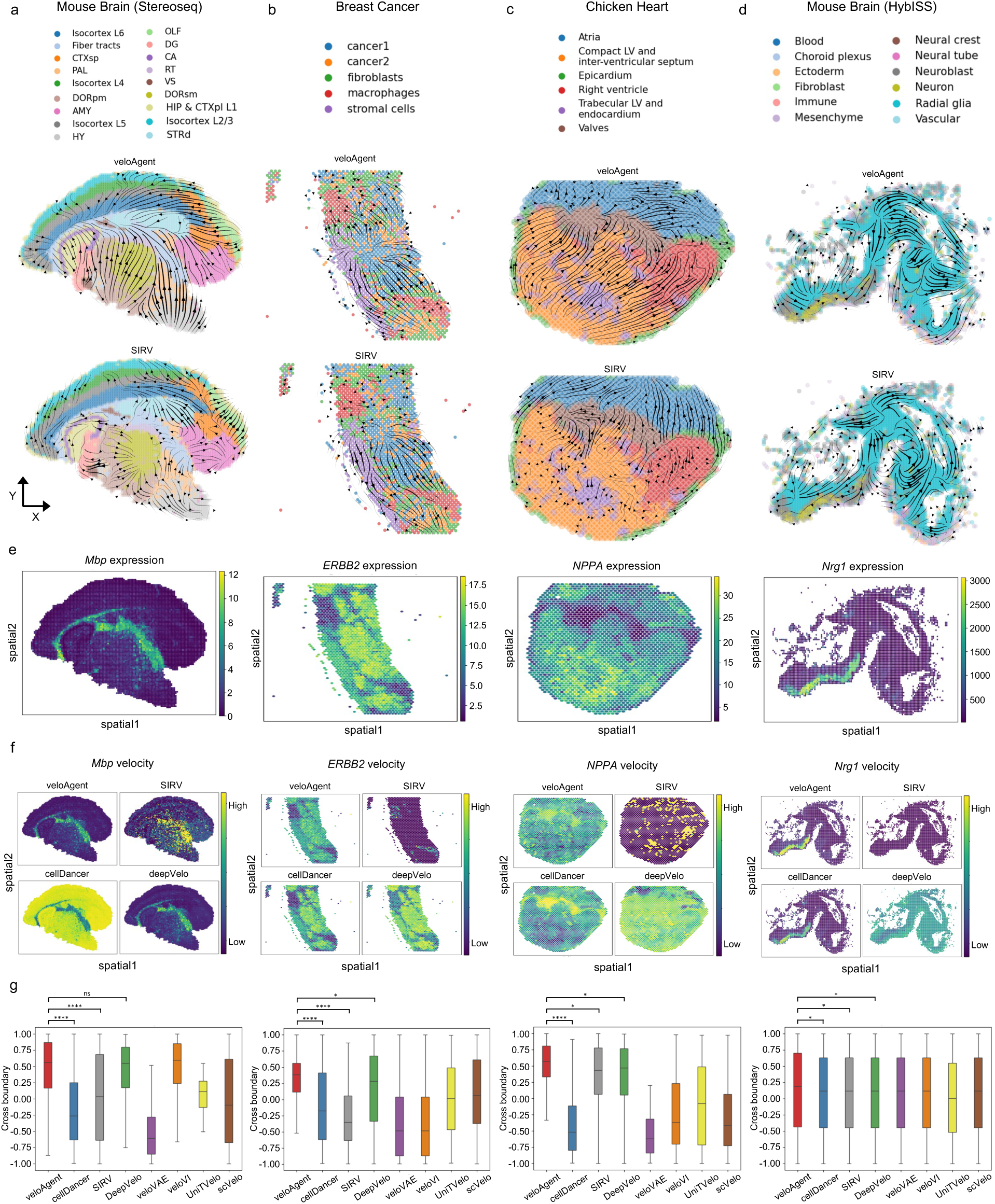
Spatial RNA velocity inference by veloAgent. veloAgent’s ability to infer and embed RNA velocity onto spatial coordinates is compared against competing methods across multiple developmental spatial transcriptomics datasets. **a-d**, Spatial RNA velocity fields computed by veloAgent (top) and SIRV (bottom), overlaid on tissue coordinate maps for: **a,** mouse brain (Stereo-seq); **b,** breast cancer; **c,** chicken heart; and **d,** mouse brain (HybISS). **e,** Spatial distribution of spliced expression for key marker genes associated with terminal cell states: *Mbp* in mouse brain (Stereo-seq)*, ERBB2* in breast cancer*; NPPA* in chicken heart*, Nrg1* in mouse brain (HybISS), visualized across spatial coordinates. **f,** Spatial RNA velocity fields for the same marker genes, comparing veloAgent (top left), SIRV (top right), cellDancer (bottom left) and DeepVelo (bottom right). veloAgent captures clearer and more localized velocity patterns toward terminal regions, consistent with expected developmental trajectories. **g,** Cross-boundary direction scores comparing veloAgent and competing methods for spatial velocity inference, measuring the correctness of predicted transitions across spatially localized cell state boundaries. Box plots show consistently higher accuracy for veloAgent. Statistical significance calculated using Mann-Whitney U test with FDR correction. Exact *P*-values can be found in Supplementary Table 3.

At the gene level, veloAgent more accurately captured the spatial dynamics of key driver genes than other methods. For terminal-state marker genes, velocity magnitudes were tightly localized to expected regions, reflecting transcriptional activation or repression near terminal states (Fig. 3e, f; Supplementary Figs. 7-8). Although expression patterns were comparable across methods, their velocity profiles differed markedly: veloAgent consistently yielded sharper and more biologically meaningful velocity patterns. For example, in the Stereo-seq mouse brain, *Mbp* velocity was concentrated within fiber tracts, consistent with its role in oligodendrocyte myelination^35^ (Fig. 3e, f; left), whereas cellDancer incorrectly predicted high velocity outside these regions. In the breast cancer dataset, *ERBB2* velocity was tightly restricted to cancer cell populations by veloAgent (Fig. 3e, f; middle left), whereas SIRV lacked this specificity despite *ERBB2* being a known oncogene driving tumor progression^36^. Similarly, in the HybISS mouse brain, *Nrg1* velocity localized to neurons in line with its role in neuronal development^38^ (Fig. 3e, f; right), while SIRV and DeepVelo produced diffuse or nonspecific signals. Even when velocity and expression were imperfectly aligned, such as for *NPPA* in the chicken heart (Fig. 3e, f; middle right), veloAgent still best reflected biology by capturing its role as an atrial differentiation marker^37^.

Quantitatively, veloAgent consistently achieved the highest scores across cross-boundary direction, spatial velocity confidence, and fate probability metrics among all datasets (Fig. 3g, Supplementary Fig 9.), confirming superior alignment between predicted velocity and spatially localized transitions. For cross boundary direction, veloAgent achieved on average 49.57% (Stereo-seq brain), 54.30% (breast cancer), 72.74% (chicken heart), and 8.75% (HybISS brain) higher accuracy than all other methods, consistently matching or surpassing competing models across datasets.

Specifically, veloAgent achieved superior performance relative to the top 3 next-best methods: Stereo-seq brain (0.5648 vs. 0.5594, veloVI; n.s) (0.5648 vs. 0.5492, DeepVelo; n.s) (0.5648 vs. - 0.1164, UniTVelo; FDR = 6.845×10^-35^), breast cancer (0.3816 vs. 0.2806, DeepVelo; FDR = 3.192×10^-2^) (0.3816 vs. 0.00654, scVelo; FDR = 2.646×10^-5^) (0.3816 vs. 0.0162, UniTVelo; FDR = 1.490×10^-9^), chicken heart (0.5744 vs. 0.4714, DeepVelo; FDR = 3.631×10^-2^) (0.5744 vs. 0.4511, SIRV; FDR = 4.325×10^-2^) (0.5744 vs -0.0759, UniTVelo; FDR = 5.793×10^-9^), and HybISS brain (0.1937 vs. 0.1273, cellDancer; FDR = 2.041×10^-2^); (0.1937 vs. 0.1228, DeepVelo; FDR = 2.041×10^-2^) (0.1937 vs. 0.1199, SIRV; FDR = 2.041×10^-2^) showing that it can consistently match or outperform other velocity inference methods.

To further assess spatial fidelity, we introduced two analog metrics—spatial velocity confidence and spatial fate probability—representing spatially mapped extensions of their conventional counterparts. On average, veloAgent achieved 46.30% (Stereo-seq brain), 13.77% (breast cancer), 29.69% (chicken heart), and 30.36% (HybISS brain) higher predictive power in spatial fate probability than competing methods (Supplementary Fig. 9b). Specifically, against the second-best method, veloAgent achieved: Stereo-seq brain (0.7250 vs. 0.5954, veloVI; FDR < 1.0×10^-308^), breast cancer (0.5648 vs. 0.5370, SIRV; FDR = 1.351×10^-2^), chicken heart (0.8331 vs. 0.7926, cellDancer; FDR = 8.071×10^-101^), and HybISS brain (0.7769 vs. 0.7280, cellDancer; FDR = 5.249×10^-40^). veloAgent demonstrated on average 36.63% (Stereo-seq brain), 36.22% (breast cancer), 33.03% (chicken heart), and 35.04% (HybISS brain) higher spatial velocity confidence than all other methods (Supplementary Fig. 9a). In direct comparison to the other models: Stereo-seq brain (0.9999 vs. 0.8576, DeepVelo; FDR = 4.406×10^-248^), breast cancer (0.9998 vs. 0.9776, cellDancer; FDR = 2.576×10^-74^), chicken heart (0.9999 vs. 0.8859, SIRV; FDR = 5.397×10^-51^), and HybISS brain (0.9999 vs. 0.9188, cellDancer; FDR = 3.157×10^-105^). veloAgent maintained superior performance across both measures, reinforcing its ability to generate accurate and spatially consistent velocity fields (Supplementary Fig. 9). Collectively, these results demonstrate that by integrating spatial priors through ABM, veloAgent transforms RNA velocity estimation from a purely temporal abstraction into a spatially coherent model of tissue dynamics, enabling more accurate and interpretable studies of cellular transitions and their underlying gene programs.

### veloAgent reveals lineage-specific driver genes and transcription factors

To determine whether the improved RNA velocity estimates from veloAgent translate into biologically meaningful gene-level insights, we leveraged CellRank as a downstream framework that operates on velocity fields^39^. Unlike prior analyses where CellRank was applied to conventional RNA velocity, here it is driven by veloAgent’s spatially and temporally coherent velocity vectors, allowing us to assess how enhanced spatial modeling improves downstream fate and driver-gene inference. For each dataset (HybISS mouse brain, chicken heart, and Stereo-seq mouse brain), we computed fate probabilities and ranked genes along pseudotime within each lineage. These ranked genes were visualized as pseudotime heatmaps, where expression was scaled and genes were ordered by the pseudotime at which smoothed expression peaked (Fig. 4a, d, g).

**Fig. 4:**
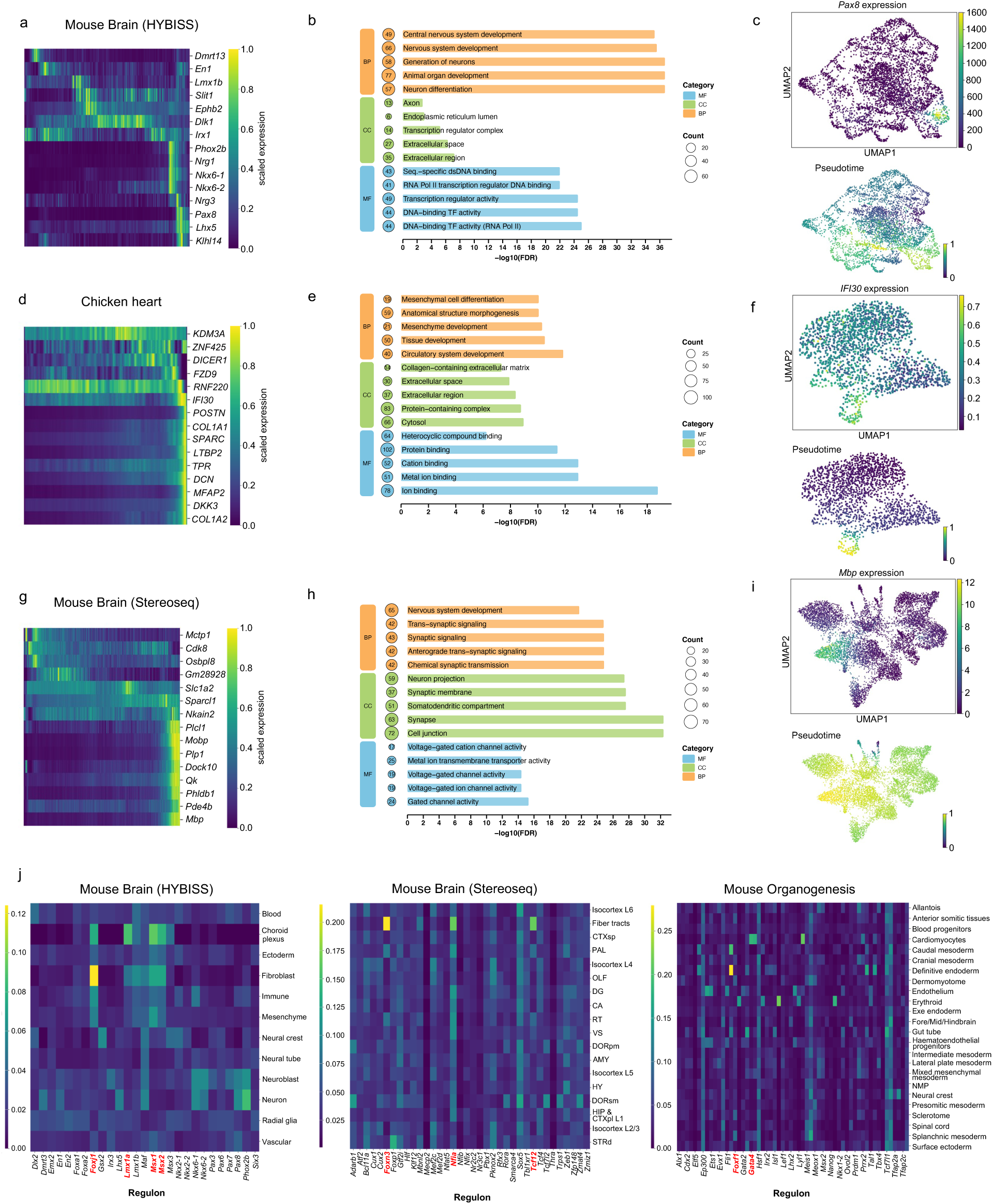
Improved driver gene and transcription factor analysis enabled by accurate RNA velocity inference from veloAgent. Accurate velocity estimates from veloAgent enhance downstream analyses such as identification of cell-type-specific driver genes and regulatory transcription factors across multiple spatial and temporal datasets. **a,** Heatmap showing normalized spliced gene expression ordered by pseudotime for driver genes identified by CellRank in neuron cell states of the HybISS mouse brain dataset. **b,** Gene Ontology (GO) enrichment for neuron driver genes from the HybISS mouse brain dataset. **c,** Spatial distribution of spliced gene expression for the top correlated driver gene, *Pax8* (top), and pseudotime projected onto the UMAP embedding (bottom). **d,** Heatmap of normalized spliced gene expression ordered by pseudotime for driver genes identified by CellRank in valve cell states of the chicken heart development dataset. **e,** GO enrichment analysis for valve driver genes in the chicken heart dataset. **f,** Spatial distribution of spliced expression for the top correlated driver gene, *IFI30* (top), and pseudotime projection on the UMAP embedding (bottom). **g,** Heatmap of normalized spliced gene expression ordered by pseudotime for driver genes identified by CellRank in fiber tract cell states of the Stereo-seq mouse brain dataset. **h,** GO enrichment analysis for fiber tract driver genes. **i,** Spatial distribution of spliced gene expression for the top correlated driver gene, *Mbp* (top), and pseudotime projection on the UMAP embedding (bottom). **j,** Heatmaps of transcription factor activity scores derived from SCENIC analysis across cell states in the mouse brain (HybISS, left), mouse brain (Stereo-seq, middle), and mouse organogenesis (right) datasets.

Across all datasets, veloAgent revealed coherent cascades of early regulators, mid-stage transition factors, and late differentiation markers that followed expected developmental orderings. In the HybISS mouse brain, veloAgent-derived pseudotime captured the sequential activation of regulators driving neuronal differentiation (Fig. 4a). Early *En1* expression aligned with progenitor-like states, consistent with its role in embryonic patterning, where loss of *En1* causes mid/hindbrain defects by E9^41^. *Dlk1* peaked at mid-pseudotime, mirroring pro-neural factors (*Ngn2*, *Mash1*, *NeuroD*) known to precede neuronal maturation^42^. Late-stage genes such as *Lhx5* and *Klhl14* were upregulated in terminal neuronal populations; *Lhx5* regulates hippocampal neuron differentiation and migration, while *Klhl14* directs corticospinal axon targeting^43–47^. These results demonstrate that veloAgent reconstructs the temporal hierarchy of transcriptional regulators underlying neurodevelopment.

In the chicken heart, veloAgent distinguished transient valve progenitors enriched for *DICER1* from terminal valve cells expressing ECM genes (*COL1A1/2*, *SPARC*) (Fig. 4d). This ordering aligns with reports that *DICER1* activity is essential for early valve development, whereas ECM expression supports structural maturation and stability of the valves later in development^48–50^. In the Stereo-seq mouse brain, veloAgent aligned early regulators such as *Cdk8*, mid-phase factors including *Quaking (Qk)*, and late markers such as *Mbp* (Fig. 4g). *Cdk8*, a Mediator-complex component, is essential for early mouse development^51,52^, while *Qk* isoforms stabilize *p27^Kip1^* mRNA to promote oligodendrocyte maturation^53^, and *Mbp* marks terminal differentiation during myelination (Fig. 4i)^54^. Together, these cascades highlight veloAgent’s ability to delineate lineage-specific transcriptional programs across distinct spatial and developmental contexts.

To assess the biological relevance of the inferred driver genes, we performed Gene Ontology (GO) enrichment analysis on the top fate-associated genes for each lineage (Fig. 4b, e, h). Enriched pathways were strongly consistent with known developmental programs. In the chicken heart (Fig. 4e), enriched terms included circulatory-system development (FDR = 1.4×10⁻¹², 40 genes), anatomical-structure morphogenesis (FDR = 8.7×10⁻¹¹, 59 genes), and mesenchymal development (FDR = 4.9×10⁻¹¹, 21 genes), implicating *POSTN* and *COL1A1* as central regulators of cardiac morphogenesis and collagen signaling^55,56^. In the HybISS mouse brain (Fig. 4b), enriched terms such as neuron differentiation (FDR = 1.9×10⁻³⁷, 57 genes), nervous-system development (FDR = 2.6×10⁻³⁶, 66 genes), and generation of neurons (FDR = 1.9×10⁻³⁷, 58 genes) confirmed neural-lineage specificity, featuring *Lhx5* and *Phox2b* as canonical regulators^47,57^. In the Stereo-seq brain (Fig. 4h), enriched pathways included nervous-system development and synaptic signaling, with *Qk* and *Mbp* emerging as key regulators of oligodendrocyte maturation and myelin formation^53,54^. These enrichment patterns confirm that the driver genes inferred from veloAgent-based velocity recapitulate lineage-regulatory programs across tissues.

To further examine upstream transcriptional regulation, we applied SCENIC^58^ to infer gene-regulatory networks and identify active transcription factors (TFs) across the HybISS, Stereo-seq, and mouse-organogenesis datasets (Fig. 4j). Because SCENIC’s predictions depend directly on the quality of the input expression and velocity information, improvements in RNA velocity modeling should translate into clearer and more biologically coherent TF-activity patterns. The predicted TFs showed strong concordance with known lineage regulators. In the HybISS dataset, *Foxj1*, *Lmx1a*, *Msx1*, and *Msx2* were recovered, consistent with their established roles in choroid-plexus patterning^59,60^ (Fig. 4j; left); *Lmx1a*, for instance, directs cells toward roof-plate and choroid-plexus lineages rather than neuronal fates originating from the rhombic lip^61^. In the Stereo-seq brain, *Foxn3*, *Nfia*, and *Tcf12* were predicted (Fig. 4j; middle); *Nfia* and *Tcf12* are established regulators of fiber-tract formation, while *Foxn3*, though not previously linked, may play an indirect regulatory role and represents a potential novel TF for further investigation^62–64^. In the mouse-organogenesis dataset, *Foxf1* and *Gata4* were identified in association with the definitive endoderm and cardiomyocyte lineages, respectively (Fig. 4j; right); *Foxf1* mediates endoderm–mesoderm signaling, and *Gata4* is a key regulator of cardiac gene expression and differentiation^65,66^.

These results demonstrate that veloAgent not only infers spatially and temporally consistent cell-state dynamics but also enables biologically validated and accurate identification of lineage-specific driver genes and transcription factors across diverse tissues.

### veloAgent facilitates in-silico perturbation of RNA velocity to reveal causal regulators and therapeutic targets

A capability absent in all existing RNA velocity inference methods—including scVelo, cellDancer, and DeepVelo—is the ability to perform *in-silico perturbations* directly within the learned dynamical system. veloAgent fills this gap by explicitly parameterizing gene-specific transcription (α), splicing (β), and degradation (γ) kinetics, enabling mechanistic simulation of how gene-level perturbations propagate through the cellular velocity field. By computationally silencing a gene’s transcription rate, veloAgent emulates a virtual knockout and recomputes the resulting flow of cell states, predicting how the loss of that gene would redirect developmental or disease trajectories in real tissue space. This approach transforms RNA velocity from a descriptive framework into a causal, predictive system capable of identifying regulatory drivers and therapeutic targets.

Perturbations were performed by systematically setting each gene’s transcription rate (α) to zero in the trained model, without retraining, and propagating the change through the dynamical equations to recalculate RNA velocity for all cells (Fig. 5a). Because the model learns gene-specific transcription, splicing, and degradation kinetics as coupled parameters, altering α post-training preserves the learned regulatory dependencies and propagates mechanistically through the velocity field rather than numerically rescaling expression. The resulting perturbed velocity fields were projected onto the UMAP embedding to visualize directional shifts. To quantify the magnitude and orientation of these shifts, we calculated the cross-boundary direction (CBD) score, which measures the change in flow vectors across boundaries separating progenitor and terminal states. A positive score indicates that silencing a gene pushes trajectories toward terminal fates, whereas a negative score reflects diversion away from them. Genes with high absolute CBD values thus act as key regulators of fate progression.

**Fig. 5.**
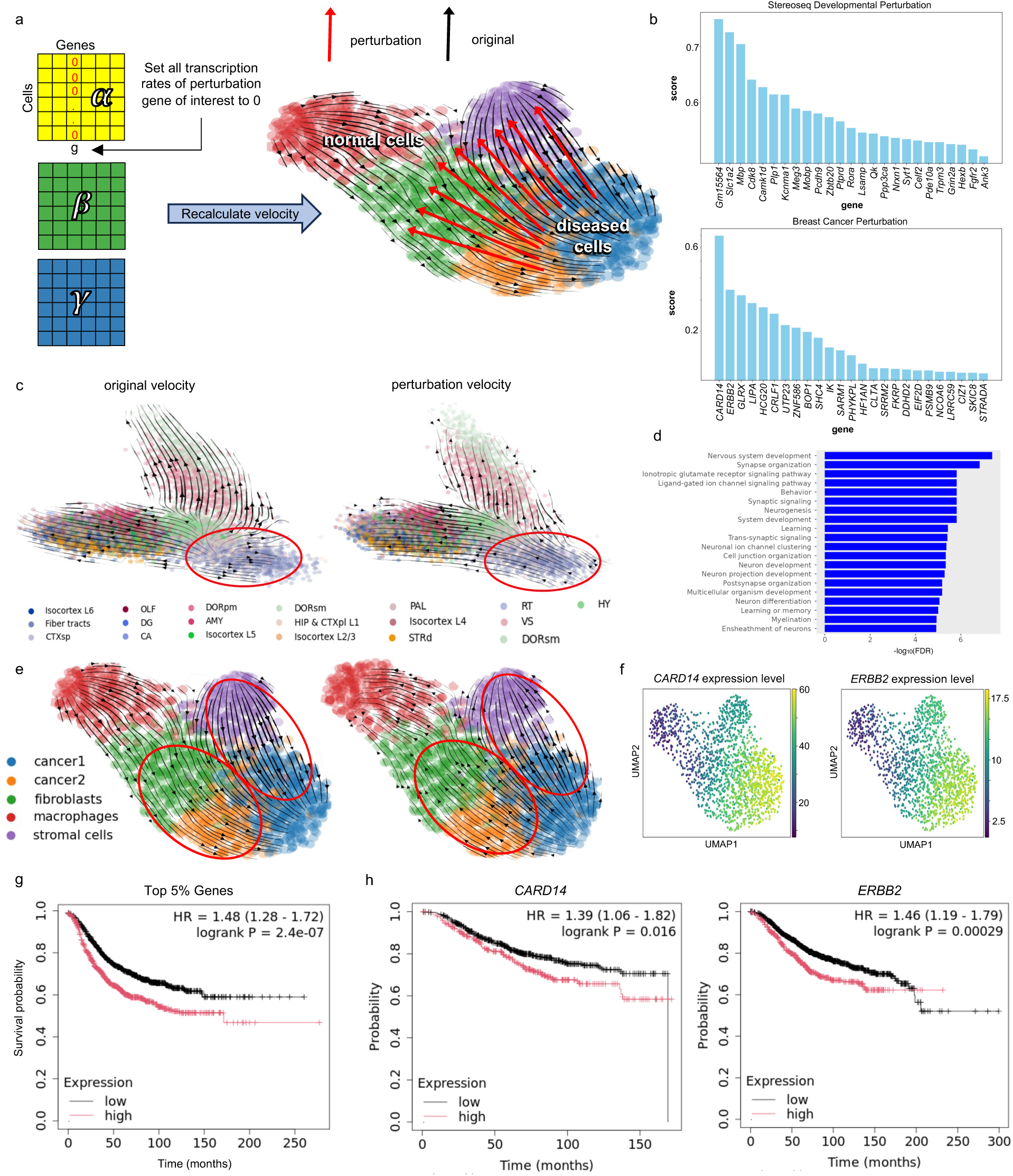
In-silico perturbation analysis enabled by veloAgent identifies potential therapeutic targets. veloAgent enables in-silico perturbation by systematically silencing transcription to assess the impact of gene-specific regulatory loss on cellular dynamics. **a,** Schematic of the perturbation pipeline, where the transcription rate (𝛼) of each gene is individually set to zero to simulate gene knockout or drug-induced repression, followed by RNA velocity recalculation. **b,** Top 5% of perturbed genes ranked by the change in cross-boundary score of the velocity following perturbation in mouse brain developmental data (top) and human breast cancer data (bottom). **c,** Comparison of perturbed versus original velocity projections in the developmental dataset, with red circles highlighting regions of substantial deviation. **d,** GO enrichment analysis of the top 5% most perturbed genes, revealing biological processes associated with developmental regulation. **e,** Velocity projection comparison in the breast cancer dataset showing notable shifts following transcriptional perturbation. **f,** UMAP projections showing expression of candidate therapeutic targets, *CARD14* and *ERBB2,* each enriched in cancer-associated cell populations. **g,** Survival analysis comparing patient outcomes between high- and low-expression groups for the combined set of significantly perturbed genes in the breast cancer dataset. **h,** Individual Kaplan-Meier survival analyses for *CARD14* and *ERBB2*, demonstrating prognostic relevance based on expression level.

We first applied this framework to a developmental mouse brain dataset profiled by Stereo-seq. The unperturbed velocity field showed expected progression toward fiber-tract cells. Silencing top-ranked genes (Fig. 5b; top), identified by high |CBD| values, reversed this directionality, implying their causal roles in guiding differentiation toward the fiber-tract fate (Fig. 5c). Gene Ontology enrichment of the top perturbed genes revealed significant over-representation of neurogenesis, axonogenesis, and synaptic-organization pathways (Fig. 5d), confirming that the perturbation-derived regulators align with known developmental programs.

We next tested whether this causal-perturbation framework could uncover regulatory dependencies in disease. Using a Visium human breast cancer spatial-transcriptomics dataset, the unperturbed velocity field indicated trajectories converging toward malignant clusters. Perturbing top-scoring genes markedly redirected the velocity vectors away from these clusters (Fig. 5e, Supplementary Fig. 10), suggesting suppression of malignant progression. The most strongly perturbed genes (Fig. 5b, bottom) included *CARD14* and *ERBB2*^67,68^—well-established oncogenes whose inhibition reversed trajectory directionality. Spatial maps confirmed their enrichment within malignant cell populations (Fig. 5f). Kaplan-Meier survival analyses showed that patients with high expression of these genes experienced significantly poorer overall survival (Fig. 5g-h). Notably, *ERBB2* is a validated therapeutic target of multiple FDA-approved drugs, including trastuzumab and trastuzumab deruxtecan^69,70^, underscoring the translational fidelity of our in-silico predictions.

To assess whether veloAgent’s perturbation framework generalizes beyond transcriptional regulation, we further simulated perturbations of the splicing rate (β) for each gene. These β-perturbations altered velocity directionality in a manner consistent with post-transcriptional control, typically inducing smoother and temporally delayed trajectory shifts relative to α-perturbations (Supplementary Fig. 11). This result demonstrates that veloAgent can model both transcriptional and splicing-driven dynamics within the same mechanistic framework, further validating its capacity for causal simulation.

By enabling gene-level kinetic perturbations within a learned velocity model, veloAgent provides a scalable, mechanistic, and hypothesis-driven alternative to physical perturbation assays. Together, these analyses establish veloAgent as a bridge between mechanistic modeling and functional discovery: by linking gene-specific kinetic parameters to emergent changes in cellular dynamics, veloAgent enables causal inference of regulatory genes, nominates candidate therapeutic targets such as *CARD14* and *ERBB2*, and offers a generalizable computational platform for virtual perturbation experiments across developmental and disease contexts.

### veloAgent is scalable and computationally robust

veloAgent’s architecture, which scales with the number of genes rather than the number of cells, provides substantial gains in computational and memory efficiency compared to existing RNA velocity models. This design choice is critical because the number of genes in each organism is essentially fixed, whereas single-cell datasets continue to expand exponentially with improvements in sequencing throughput. In contrast, most frameworks—including cellDancer—scale with cell count, leading to rapidly increasing memory and runtime requirements that limit their applicability for atlas-level analyses involving hundreds of thousands or millions of cells.

To systematically benchmark scalability, we progressively duplicated the HybISS mouse brain dataset to generate datasets ranging from 50,000 to one million cells. While cellDancer’s memory usage scaled near cubically, reaching ∼120 GB at one million cells, veloAgent exhibited near-linear scaling, requiring only ∼37 GB under identical conditions (Fig. 6a, top; Supplementary Fig. 12). Similarly, veloAgent completed velocity inference substantially faster, maintaining nearly constant per-cell runtime as dataset size increased (Fig. 6a, bottom; Supplementary Fig. 13). cellDancer was selected as the primary comparator because it is the most competitive deep-learning–based RNA velocity framework and offers a direct architectural contrast to veloAgent’s gene-scaled design. Other frameworks such as scVelo were excluded from this analysis because their smaller network architectures preclude meaningful efficiency comparisons, though they were included in biological benchmarking elsewhere in this study. Full experimental and hardware details are provided in the Methods section.

**Fig. 6.**
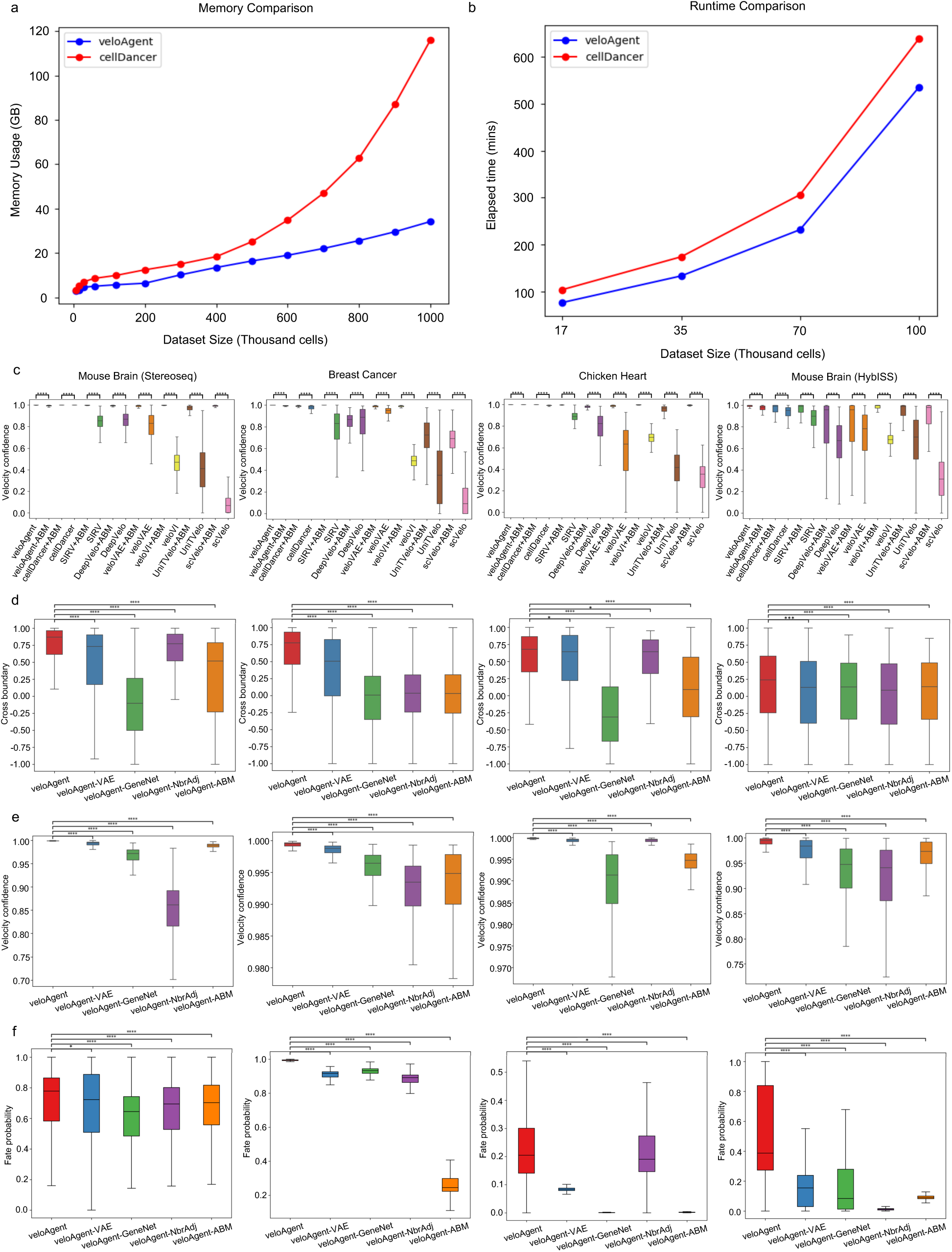
Ablation study and computational efficiency of veloAgent. **a,** Comparison between veloAgent and cellDancer regarding peak CPU memory usage (top) for increasing number of cells subsampled from HybISS mouse brain dataset. **b,** Comparison of computational runtime on CPU for increasing number of cells, subsampled from HybISS mouse brain dataset between veloAgent and cellDancer. **c,** Comparative evaluation of velocity confidence scores before and after applying veloAgent’s agent-based model (ABM) for spatial refinement on top of each method’s temporal velocity estimates. Original estimates from each method are contrasted with their spatially adjusted counterparts, showing consistent improvement when spatial data is incorporated. **d,** To assess the contribution of key model components, an ablation study was performed by systematically removing each module from veloAgent. Cross-boundary accuracy, velocity confidence and fate probability metrics across four configurations: full veloAgent, without latent-space smoothing, without gene-informed network architecture, without neighbour adjustment and without incorporating spatial transcriptomics data using agent-based modeling. Performance drops observed in ablated models validate the importance of each component. Statistical significance calculated using Mann-Whitney U test with FDR correction. Exact *P*-values can be found in Supplementary Table 4.

Beyond computational scalability, veloAgent enhances biological robustness through its agent-based modeling (ABM) module, which integrates spatial context to refine velocity fields. The ABM takes temporally inferred velocity estimates and couples them with spatial coordinates of neighboring cells to produce spatially coherent velocity directions. Notably, this module is fully modular and can be applied to improve the outputs of other RNA velocity methods without retraining. Across all datasets, ABM integration markedly increased median velocity confidence relative to scVelo (Fig. 6c): mouse brain (Stereo-seq, 0.9943 vs. 0.0677; FDR < 1.0 × 10⁻³⁰⁸), breast cancer (0.6920 vs. 0.0929; FDR < 1.0 × 10⁻³⁰⁸), chicken heart (0.9959 vs. 0.3550; FDR < 1.0 × 10⁻³⁰⁸), and mouse brain (HybISS, 0.9794 vs. 0.3166; FDR < 1.0 × 10⁻³⁰⁸). These consistent gains demonstrate that the ABM functions as a general-purpose enhancement, improving spatial coherence while preserving veloAgent’s computational efficiency.

To evaluate model robustness, we conducted ablation experiments on spatial transcriptomics datasets using four complementary metrics: cross-boundary direction, fate probability, velocity confidence, and in-cluster coherence. Removing any of veloAgent’s key components—the VAE encoder (Phase 1), gene-gene interaction network (Phase 2), neighborhood adjustment (Phase 3), or ABM refinement (Phase 4)—significantly reduced performance, whereas the complete model consistently achieved the highest accuracy (Fig. 6d-f; Supplementary Fig. 14). Incorporating spatial context through the ABM provided the greatest improvement, underscoring its importance for biologically coherent velocity estimation. Ablation analyses on scRNA-seq-only datasets (without ABM) yielded consistent results (Supplementary Fig. 15).

## DISCUSSION

In this study, we present veloAgent, a deep generative and agent-based model (ABM) for spatial RNA velocity estimation that incorporates spatial transcriptomics data. Most existing RNA velocity frameworks treat cells as independent points in expression space, neglecting the tissue architecture and interactions between cells that influence cell state transitions. veloAgent bridges this gap by combining a VAE for robust latent representation, a gene-regulatory-aware neural network that incorporates known molecular interactions, and an ABM that simulates each cell’s spatial microenvironment. This integration ensures that velocity estimates are not only informed by transcriptional dynamics but also constrained by spatial neighbourhood structure, improving both accuracy and interpretability. Applied to developmental and disease datasets, veloAgent outperforms existing methods across quantitative benchmarks and captures literature-supported marker gene dynamics in their correct spatial contexts. Beyond trajectory inference, veloAgent’s in silico perturbation module predicts how changes in gene regulation or signaling alter cell fate, enabling targeted hypothesis generation in developmental biology oncology and regenerative medicine.

veloAgent offers several key advances over existing RNA velocity frameworks. **First**, it is designed for scalability through a gene-centric architecture, where model complexity scale with the number of genes —typically fixed for an organism— rather than the number of cells. This contrasts with cell-centric methods such as cellDancer, whose complexity grows with cell count, making large-scale analysis computationally expensive. By scaling with genes instead of cells, veloAgent achieves sublinear memory usage and can handle atlas-scale datasets with millions of cells. This is particularly important given the recent rise in single-cell experimental throughput, which is expected to continue growing. Unlike models that rely on global assumptions or cell-centric designs, our approach generalizes across tissues and experimental platforms without compromising interpretability or performance^71^. **Second**, veloAgent uniquely integrates spatial transcriptomics data into the velocity inference process through its ABM module. Spatial context is often overlooked in RNA velocity frameworks, yet it plays a fundamental role in shaping lineage boundaries, localized gene expression patterns, and environmental signals that guide cell behavior^72,73^. By simulating the spatial microenvironments of each cell, veloAgent embeds tissue architecture into its RNA velocity estimations, ensuring that transcriptionally similar and spatially proximal cells are inferred to follow coherent trajectories. In contrast, existing approaches either omit spatial information entirely or use it only for post hoc projection (e.g. SIRV), which can visualize velocities on a tissue map but cannot influence the underlying estimation. veloAgent unifies both, incorporating spatial constraints during velocity inference and projecting the resulting vectors back into physical space to reveal region-specific linear trajectories and niche-dependent differentiation. **Third**, veloAgent’s modular ABM design allows it to function as a universal booster, seamlessly integrating with existing RNA velocity frameworks. By coupling their outputs with spatial context, the ABM refines and harmonizes velocity fields across tissue regions, substantially enhancing spatial coherence and biological interpretability without retraining of the original models. **Finally**, beyond trajectory inference, veloAgent introduces an in-silico perturbation framework that allows targeted manipulation of RNA velocity vectors to simulate the effects of gene regulatory or signalling interventions. This enables rational cell state steering and identification of candidate therapeutic targets. As a case study, we show that veloAgent can recover key breast cancer linked genes whose perturbation could reverse disease trajectories, including *ERBB2*, which has existing FDA-approved inhibitors. This opens new avenues for therapeutic discovery, such as identifying candidate drug targets or designing intervention strategies for regenerative medicine and cell reprogramming.

While veloAgent offers significant advancements, it also faces limitations that warrant future investigation. The current perturbation framework models single gene interventions for simplicity and does not capture the complexity of combinatorial perturbations or the interplay of multiple regulatory pathways. Additionally, integrating other omics modalities such as ATAC-seq and proteomics could further enhance both velocity estimation as well as perturbation modeling by providing a more complete view of cell state and regulatory control. Finally, while veloAgent is designed for scalability, it has not yet been applied to a true atlas dataset with millions of cells, representing an important step to fully demonstrate its large-scale applicability.

veloAgent defines a scalable, spatially aware, and biologically interpretable framework that extends RNA-velocity analysis into previously inaccessible domains. Its scalability supports emerging cell-atlas projects, and its perturbation module bridges computational modelling with experimental design. The framework provides a foundation for predictive, intervention-ready models of cellular behaviour, with the potential to accelerate discovery, advance developmental understanding, and guide therapeutic innovation.

## METHODS

### Single-cell sequencing data preprocessing

We processed the single-cell RNA-sequencing data using the scVelo pipeline. We began by filtering out cells with less than 20 genes. We then selected the top 2,000 highly variable genes and normalized against the library size and transformed into log space. We then computed a nearest-neighbor graph (with 30 neighbors) in principal component analysis space (30 principal components). Finally, we calculated the first-order moment of spliced and unspliced counts for subsequent velocity estimation steps.

### veloAgent framework

The veloAgent model consist of three sub-components, a variational autoencoder (VAE), a deep neural network, and an agent-based model (ABM).

#### Phase 1: Variational Autoencoder

veloAgent uses a VAE designed to represent high-dimensional single-cell gene expression data within a lower-dimensional embedding space. Specifically, our VAE incorporates two separate encoders, each sharing an identical structure composed of three fully connected hidden layers. These hidden layers progressively decrease in dimension, evenly spaced in thirds, ultimately reducing the input to the latent dimension 𝑍. Each encoder, 𝑞*_θ_*, independently process the spliced 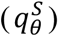 and unspliced 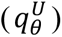 gene expression, transforming them into distinct low dimensional embeddings. The outputs from both encoders are concatenated to form a unified representation, 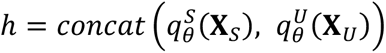. Two MLPs are then employed to determine the mean vectors, 𝜇*_z_* = 𝑓_𝜇_*θ*__(𝜇*_z_*|ℎ), and standard deviation vectors 𝜎*_z_* = 𝑓_𝜎_*θ*__(𝜎*_z_*|ℎ) of the latent representation, assuming a multivariate normal distribution as a prior, 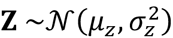. The approximated posterior distribution is represented as 𝑞*_θ_*(𝐙 | 𝐗*_S_*, 𝐗*_U_*). The latent representation is particularly valuable, as they are designed to preserve critical biological information while reducing noise and less relevant features. This enables more coherent and accurate estimations of single-cell velocities. Two corresponding decoders 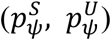 mirror the encoders architectures reconstructing the original cell-by-gene matrices for splices and unspliced counts from 𝐙. The VAE loss function is given by the following equation:

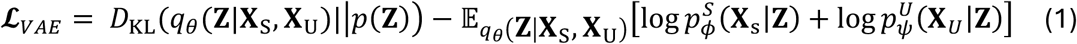

#### Phase 2: Velocity Rate Estimation using Deep Neural Network

The transcriptional kinetics of an individual gene, 𝑔, are modeled by a system of two ordinary differential equations:

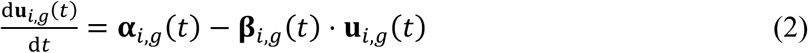

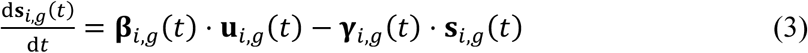

where 𝐮*_i,g_* (𝑡) and 𝐬*_i,g_* (𝑡) represent the concentrations of the premature and mature mRNAs for cell 𝑖, gene 𝑔 at time 𝑡 and 𝛂*_i,g_*, 𝛃*_i,g_*, 𝛄*_i,g_* represent the transcription, splicing and degradation rates at time 𝑡, respectively. Previous methods assume that parameters **α**, **β**, and **γ** are either constant or universally shared across all genes and cells. However, this assumption inadequately models the continuously evolving and heterogeneous cell subpopulations.

To accurately estimate **α**, **β**, and **γ**, we utilize a neural network that takes as input the latent representation 𝐙 generated by the VAE in phase 1. The network initially contains a fully connected hidden layer with dense connections from the latent dimension ( Z) to the gene expression dimension (*G*). The second hidden layer incorporates sparse connections informed by gene-gene interaction data from the STRING database. Specifically, connections in this layer reflect known or predicted gene-gene interactions from STRING, including physical interactions and functional relationships inferred from co-expression, literature mining and curated databases. Genes without known connections are connected to all other gene nodes, resulting in a fully connected node for that gene. Following this biologically informed layer, the network proceeds through two additional fully connected layers, expanding first to 2*G* and then to 3*G*, where *G* is the number of genes. The final layer is structured such that the first *G* units correspond to transcription rate (**α**), the second *G* units correspond to splicing rate (**β**), and the third *G* units correspond to degradation rate (**γ**). The output provides independent rate parameters for each gene in every cell, resulting in cell and gene-specific estimates of kinetic rates.

The neural network is trained using a loss function based on cosine similarity to compare the predicted RNA velocity with observed velocity vectors. Given the outputs **α**, **β**, and **γ**, the predicted spliced (*ŝ*) and unspliced (*û*) counts are obtained using the ordinary differential equations. The velocity vector for each cell is defined as the change from its observed counts to its predicted counts:

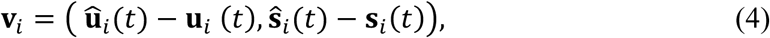

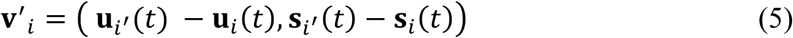

where {𝑖′} is the collection of neighboring cells of cell 𝑖; and 𝑖′ is the neighboring cell that minimizes the cosine similarity loss function. To assess the quality of the predicted velocity, we compute the cosine similarity between the velocity of cell *i,* 𝑣*_i_*, and the velocity vectors of neighboring cells:

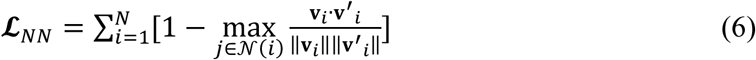

where N is the number of cells and 𝒩(𝑖) is the neighborhood of cell 𝑖, and 𝑗 is a neighboring cell of 𝑖. We take the maximum cosine similarity between the cell’s predicted velocity and the velocity of all its neighbors.

#### Phase 3: Cell Velocity Adjustment

In this phase, we refine each cell’s predicted velocity to produce a more consistent and confident estimate by leveraging information from neighbouring cells. Specifically, we adjust each cell’s velocity to align with those of its neighbours by minimizing the cosine similarity between their velocity vectors. This alignment step enhances the stability and reliability of the velocity estimates by encouraging local consistency in the velocity field. The underlying assumption is that cells within a local neighborhood in gene expression space tend to follow similar dynamic patterns. The loss function used for this alignment is formally defined below, where 𝑁 is the number of cells, 𝒩(𝑖) is the neighborhood of cell 𝑖, and 𝑗 𝜖 𝒩(𝑖) is a neighboring cell of 𝑖:

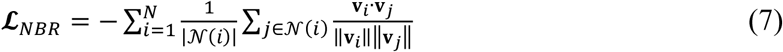

The overall loss function of veloAgent combines three components: the VAE reconstruction loss 𝓛*_VAE_*, the rate prediction loss from the neural network 𝓛*_NN_*, and the neighborhood coherence loss 𝓛*_NBR_*. Together, these form the total training objective of the model, expressed as:

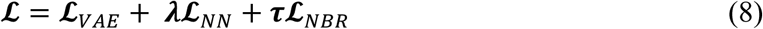

#### Agent-based Model for Spatial RNA Velocity Modeling

To incorporate spatial context into RNA velocity estimation, we developed an ABM in which each cell’s velocity is influenced by its neighboring cells within a defined spot radius. ABMs have previously been used to model cellular dynamics in single-cell and spatial transcriptomics data, capturing complex interactions among cells in a spatially aware manner. The model refines velocity estimates based on three key factors: local cell density, inverse distance, and RNA expression similarity. To construct the ABM, we first define the states and behaviours of the agent, where each agent represents a cell in the biological system. Agent states capture the properties and dynamic features of each cell, while agent rules define how each agent behaves.

##### Cell Agent State Description

Since our primary goal is to study RNA velocity in the context of spatially organized tissues, each cell agent is characterized by features relevant of both its transcriptional state and its spatial interactions. These include gene expression profiles, RNA velocity vectors, and spatial coordinates. We also include metadata such as cell type or tissue annotations, batch ID etc.

##### RNA Velocity Adjustment Behaviour Rules

The **agent behaviors** are defined by rules governing how each cell’s velocity is modulated by its neighbors. For a given cell 𝑖, the neighborhood 𝑁(𝑖) comprises all cells located within a specified radius 𝑟. The updated velocity of cell 𝑖 is computed as a weighted combination of its own velocity and those of its neighbors, reflecting three biologically motivated influence terms. First, neighboring cells with similar expression profiles are expected to exhibit similar transcriptional dynamics. This relationship is captured by a similarity weight defined as

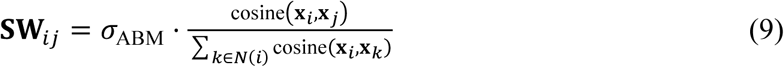

Here, 𝜎_ABM_ is a scaling constant.

Second, denser local environments exert greater collective influence on a cell’s velocity, quantified by a density weight

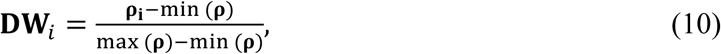

Third, spatial proximity modulates intercellular influence, as nearby cells are more likely to affect one another through diffusion-limited molecular signaling that alters gene expression. This effect is modeled using inverse-distance weighting, assigning stronger influence on closer neighbors.

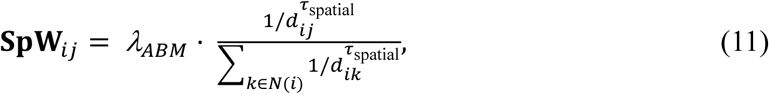

where 𝑑*_ij_* is the Euclidean distance between cells 𝑖and 𝑗, and 𝜏_spatial_ = 2, for 2D data.

The refined velocity for each cell 𝑖 is then obtained by combining its intrinsic and neighborhood influences according to

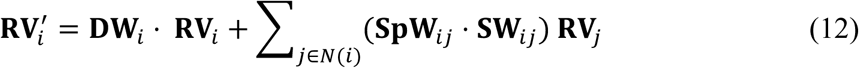

This formulation ensures that transcriptionally similar and spatially proximal cells exhibit coherent directional dynamics, while preserving biological heterogeneity across distinct tissue regions. Through this spatial refinement, the ABM transforms RNA velocity into a tissue-consistent and biologically interpretable model of cellular state transitions.

### In silico Perturbation

To evaluate the contribution of individual genes to RNA velocity and trajectory inference, we implemented an *in silico* perturbation framework within veloAgent. For each gene, a virtual loss-of-function was simulated by setting its transcription rate parameter (α) to zero, effectively removing its transcriptional influence from the learned kinetic system. The model was then used to recompute RNA velocity estimates (𝑣^(–*g*)^) without retraining, allowing direct assessment of how the perturbation altered predicted cellular dynamics.

The impact of each perturbation was quantified using the **cross-boundary direction (CBD)** metric. This metric quantifies how well velocity vectors maintain the correct directionality between transitioning cell types (e.g. cell type A to cell type B), reflecting the gene’s influence (𝐈*_g_*) on the inferred dynamic processes CBD(𝐯, 𝑑(𝐴, 𝐵)). The change in CBD following perturbation, CBD(𝑣^(-*g*)^, 𝑑(𝐵, 𝐴)), reflects the influence of gene *g* on the inferred dynamic process. A gene-level importance score was computed as

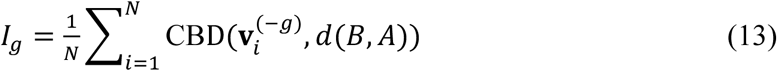

where 𝑁 denotes the total number of cells. This score represents the extent to which perturbing a specific gene alters trajectory directionality, thus identifying regulators that drive or constrain lineage progression. Statistical significance was determined using permutation testing to identify genes whose perturbation caused a significant disruption in velocity alignment.

To assess the biological relevance of these computational perturbations, we performed Reactome pathway enrichment analysis for the mouse brain developmental dataset, confirming that top-ranked perturbed genes were enriched for neurogenesis- and differentiation-related pathways. For the breast cancer dataset, we validated candidate therapeutic genes by examining their association with patient survival using Kaplan-Meier (KM) analysis, linking predicted regulatory effects to clinical outcomes.

### Benchmarking

We benchmarked veloAgent against seven state-of-the-art RNA velocity frameworks encompassing both conventional and spatially aware approaches. Among these, SIRV is the only method that also uses spatial transcriptomics data for spatial velocity projection. Comparisons were performed on scRNA-seq datasets for all methods except SIRV and on spatial transcriptomics datasets for all methods.

Performance was evaluated using three established metrics: **velocity confidence**, **cross-boundary direction**, and **fate probability accuracy**. Velocity confidence measures the local cosine similarity between a cell’s velocity vector and those of its neighbors, reflecting the global coherence and reliability of the inferred velocity field. Cross-boundary direction assesses whether velocity vectors point in the biologically expected direction across annotated lineage transitions, quantifying directional accuracy at cell-type boundaries. Fate probability accuracy evaluates how well each method captures the commitment of cells toward terminal states based on their inferred RNA velocities. Fate probabilities were computed by constructing velocity-based transition matrices and quantifying the agreement between inferred and known terminal states.

### Ablation

To evaluate the contribution of each major component in the veloAgent framework, we conducted systematic ablation experiments in which individual modules were selectively removed or simplified. Four variants were tested: the VAE, the gene-gene interaction network, the neighbor refinement mechanism, and the agent-based model (ABM). In the veloAgent-VAE variant, the learned VAE was replaced with a fixed principal component analysis (PCA) projection of the spliced and unspliced counts, eliminating the ability to learn a nonlinear latent space that captures complex cellular structure. In the veloAgent-GeneNet variant, the biologically informed gene-gene interaction layer was substituted with a fully connected layer, removing the inductive bias derived from known molecular relationships and thereby reducing the biological interpretability of the learned dependencies. The veloAgent-NBR variant ablated the neighborhood refinement step, which aligns velocity vectors across transcriptionally similar cells to ensure local directional consistency. The veloAgent-ABM variant excluded spatial refinement, preventing the model from leveraging spatial transcriptomics data to enhance tissue-level coherence and directional accuracy. All ablation variants were evaluated using the same three metrics—cross-boundary correctness, velocity confidence, and fate probability—to quantify the contribution of each component.

### Driver Gene Analysis

To identify key driver genes underlying cellular differentiation trajectories, we performed a pseudotime analysis using RNA velocity-based ordering. After computing the velocity graph, we derived pseudotime estimates for each cell using scv.tl.velocity_pseudotime. We then visualized the expression dynamics of genes of interest (GOIs) across pseudotime using a heatmap. The selected GOIs were identified from CellRank lineage analysis as putative drivers of specific fate transitions. The heatmap was sorted by velocity pseudotime to capture temporal expression trends, highlighting the sequential activation and repression of lineage-associated genes along differentiation paths.

To further validate the relevance of lineage-driving genes identified through CellRank, we extracted all genes significantly associated with a specific terminal fate based on their correlation with fate probabilities. We then performed Gene Ontology (GO) enrichment analysis on this set of genes to evaluate the biological processes and pathways most enriched within each fate, thereby confirming the functional relevance of the identified gene programs.

### Transcription Factor Analysis

To investigate the transcriptional regulators associated with terminal fates, we performed transcription factor (TF) inference using SCENIC pipeline. Transcription factor-target gene relationships were first inferred from gene co-expression patterns and then refined by assessing the enrichment of transcription factor binding motifs in the regulatory regions of target genes, using a curated database of mouse DNA motifs derived from experimental evidence. Regulon activity was quantified using AUCell and summarized by cell type to reveal distinct regulatory programs associated with specific cellular identities.

### Computer Configurations

Computational experiments were conducted on a high-performance computing server running Ubuntu 20.04.6 LTS with Linux kernel 5.4.0-216-generic (x86_64). The system was equipped with two Intel® Xeon® Platinum 8160 CPUs (96 logical cores, 2.10 GHz), 500 GB of RAM, and 8 GPUs (four NVIDIA Tesla M10 GPUs, 8 GB each; and four NVIDIA A16 GPUs, 15 GB each).

## DATA AVAILABLITY

All scRNA-seq data in this study were downloaded publicly. For the pancreatic endocrinogenesis and erythroid lineage of the mouse gastrulation datasets, we retrieve them from the Bergen et al. in the scVelo^10^. The mouse hippocampal dentate gyrus neurogenesis data was obtained from the study by La Manno et al^9^. The HybISS developing mouse brain, developing chicken heart, and SeqFISH mouse organogenesis spatial datasets were taken from the SIRV study by Abdelaal et al ^12^. The Stereo-seq mouse brain data was obtained from STOmicsDB publicly available dataset collection from the study by Chen, Ao et. al ^26^. The human breast cancer data was obtained from 10X genomic published datasets^74^.

## CODE AVAILABLITY

veloAgent is implemented in Python and available at https://github.com/mcgilldinglab/veloAgent.

## ACKNOWLEDGEMENTS

This work is supported by grants from the Canadian Institutes of Health Research (CIHR) [PJT-180505 to J.D.]; the Funds de recherche du Québec—Santé (FRQS) [295298 to J.D., 295299 to J.D., 366764 to J.D.]; the Natural Sciences and Engineering Research Council of Canada (NSERC) [RGPIN2022-04399 to J.D.]; and the Meakins-Christie Chair in Respiratory Research [to J.D.]; V.R received support from the Canada First Research Excellence Fund and the Fonds de recherche du Quebec awarded to the D2R Initiative at McGill University. This research was enabled in part by support provided by Calcul Quebec (calculquebec.ca) and the Digital Research Alliance of Canada (alliancecan.ca).

